# Pan-proteome regulation of marine *Synechococcus* nutrient stress plasticity

**DOI:** 10.64898/2026.09.21.753132

**Authors:** NS Garcia, M Saito, R Harcourt, Matthew R. McIlvin, AC Martiny

## Abstract

Variable phytoplankton resource demands can buffer productivity to climate-driven declines in nutrient supply, but the extent of this buffering depends on limits to cellular elemental plasticity. Elemental plasticity is driven by a combination of genomic adaptation and physiological regulation, but their mechanistic links remain unresolved. Here, we combine continuous culture experiments with global proteomics and elemental analyses to determine how genomic diversity shapes nutrient stress plasticity in the globally distributed cyanobacterium *Synechococcus*. By maintaining fixed growth rates in chemostats under nitrogen- or phosphorus-limiting conditions, we disentangle lineage-specific resource demands from growth effects. All strains show conserved induction of nutrient acquisition pathways, yet strain identity explains most pan-proteome variance, and variation in non-core protein expression predicts deviations in elemental plasticity. Strains from oligotrophic compared to mesotrophic regions exhibit broader proteomic adjustments and wider C:P and N:P ranges. Our findings show that genomic diversity, expressed through pan-proteome regulation, amplifies physiological plasticity and increases phytoplankton nutrient stress buffering capacity under future climate conditions.

**Importance:** Marine cyanobacteria are among the most abundant microbes on Earth and play central roles in global nutrient cycling, yet the molecular mechanisms that determine how they respond to nutrient stress remain poorly understood. Here we show that genomic diversity within the marine cyanobacterium *Synechococcus* translates into differences in protein expression and cellular resource allocation. Combining chemostat experiments and proteomics across multiple strains, we demonstrate that variation in the non-core genome shapes how lineages regulate nutrient acquisition pathways and adjust cellular C:N:P stoichiometry under nitrogen or phosphorus stress. These results reveal how pan-genome diversity generates physiological plasticity within microbial populations, providing a mechanistic link between microbial genomic variation and ecosystem-level nutrient cycling in the ocean.

## Introduction

The elemental composition and stoichiometry of carbon-to-nutrients in phytoplankton (i.e., cellular C:N:P) are fundamental features of ocean ecosystems (1). Ecologically, the elemental composition affects the quality of phytoplankton as a food source for grazers. Biogeochemically, many ocean ecosystems operate in ways that mimic a continuous culturing system or chemostat. Ocean stratification regulates the inflow of nutrients that are almost completely assimilated. Hence, the carbon-to-nutrient ratio controls the resulting carbon-based productivity and biomass stocks given a limiting nutrient flux. Future ocean warming and stratification may severely restrict the vertical flux of nutrients and hence suppress productivity (2). However, the elemental composition and associated nutrient requirements (i.e., cellular C:N:P) can vary among phytoplankton (1, 3). Consequently, flexibility in phytoplankton nutrient demand can buffer biomass and productivity to climate-driven stratification and a declining nutrient supply (4, 5). The strength of this ‘buffering’ depends on the degree of cellular elemental plasticity and hence limits to their nutrient requirements to sustain growth. Modeling studies predict that future net primary production could decline sharply if phytoplankton elemental composition is fixed (no plasticity nor buffering), remain stable with moderate plasticity (weak buffering), or even increase if elemental requirements are highly plastic (strong buffering) (5). Yet, the physiological and evolutionary factors that define the boundaries of tolerable resource stress remain poorly constrained, even though these limits are likely critical in shaping future ecosystem functions and biogeochemical feedbacks to climate change.

Phytoplankton responses to nutrient stress are governed by a combination of evolutionary adaptation and physiological regulation. Many lineages, including the globally abundant cyanobacterium *Synechococcus*, have a core set of genes present in all members, but differ markedly in non-core gene content (i.e., genome difference among lineages), forming a large pan-genome (6, 7). This genomic diversity underpins ecological diversification across environmental gradients and enables Cyanobacteria to occupy diverse marine biomes (8, 9). Gene gains and losses are recurrent signatures of adaptation to low-nutrient environments and specific elemental stresses such as nitrogen or phosphorus scarcity (10, 11). Moreover, variable presence of non-core genes involved in nutrient acquisition and storage correlates in nature with shifts in cellular elemental demands and C:N:P stoichiometry (12, 13). However, most links between genomic variation and biogeochemical outcomes remain correlative (14). We still lack a mechanistic understanding of how adaptation and associated genome diversity shape the biochemical pathways that determine cellular resource demands and cellular C:N:P.

In contrast to the long-term genetic adaptation that generates substantial genomic diversity among Cyanobacteria, physiological regulation governs how individual cells respond to immediate environmental change. Numerous studies show that phytoplankton exhibit strong and coordinated physiological responses to nutrient stress, including changes in expression of growth, nutrient acquisition, and storage pathways (11, 15, 16). These cellular adjustments define trait plasticity, namely the capacity of a lineage to redistribute metabolic and structural investments as nutrient supply changes. This physiological regulation sets the limits of resource imbalance that a lineage can tolerate and thus constrains elemental quotas and limits to C:N:P ratios (17). While genomic comparisons reveal extensive variation in metabolic potential among lineages, and physiological studies demonstrate pronounced plastic responses, the mechanistic link between pan-genome diversity and pan-proteome regulation remains unresolved. This link is poorly understood not only in *Synechococcus* but also in other phytoplankton and bacteria in general (18). As a result, we still lack a clear understanding of how variations in genome and proteome translate into the realized limits of elemental plasticity.

Here, we test how genomic diversity shapes proteomic composition - the *pan-proteome* - and resulting elemental stoichiometry within *Synechococcus*. Interpretations of elemental plasticity are often confounded by growth-rate effects, because cells can achieve extreme C:N:P ratios under slow growth (1, 19). We use continuous culture (chemostat) experiments to maintain a fixed growth rate across genetically diverse strains under nitrogen- or phosphorus-limiting conditions. Combining data-independent acquisition mass spectrometry with cellular elemental analyses (15, 20), we then investigate how genome variation governs nutrient stress responses and resource demands. By contrasting strains adapted to distinct nutrient regimes, we reveal how evolutionary adaptation and physiological plasticity interact to structure the *Synechococcus* pan-proteome, nutrient stress response, and its biogeochemical role in the ocean.

## Results

To determine the role of genomic variation for the regulation of trait expression and cellular elemental composition, we studied six marine strains within the abundant genus *Synechococcus* (Fig. 1 and S1). The strains are picked to represent major *Synechococcus* clades, spanning both mesotrophic (clade I and IV) and oligotrophic (clade II and III) representatives (Table 1) (21). The strains are closely related, but the average whole genome amino-acid identities (AAI) and shared gene content range between the strains (Fig. S1). The core genome across these strains consists of 1822 genes and covers between 52% (strain ROS8604_meso_) and 77% (CC9902_meso_) of the overall genome (Fig. 1A). The non-core genome varies among the strains (Fig. 1). For example, some genes related to nitrogen and phosphorus are core but others are variable among the strains (Fig. 1B, C), showing a difference in genome content also seen among populations of *Prochlorococcus* and *Synechococcus* (13, 22). Generally, the gene content and trait similarity do not relate strongly with phylogenetic clade membership (Fig. S1), suggesting a non-phylogenetic distribution of non-core genes and key nutrient stress traits.

**Figure 1.**
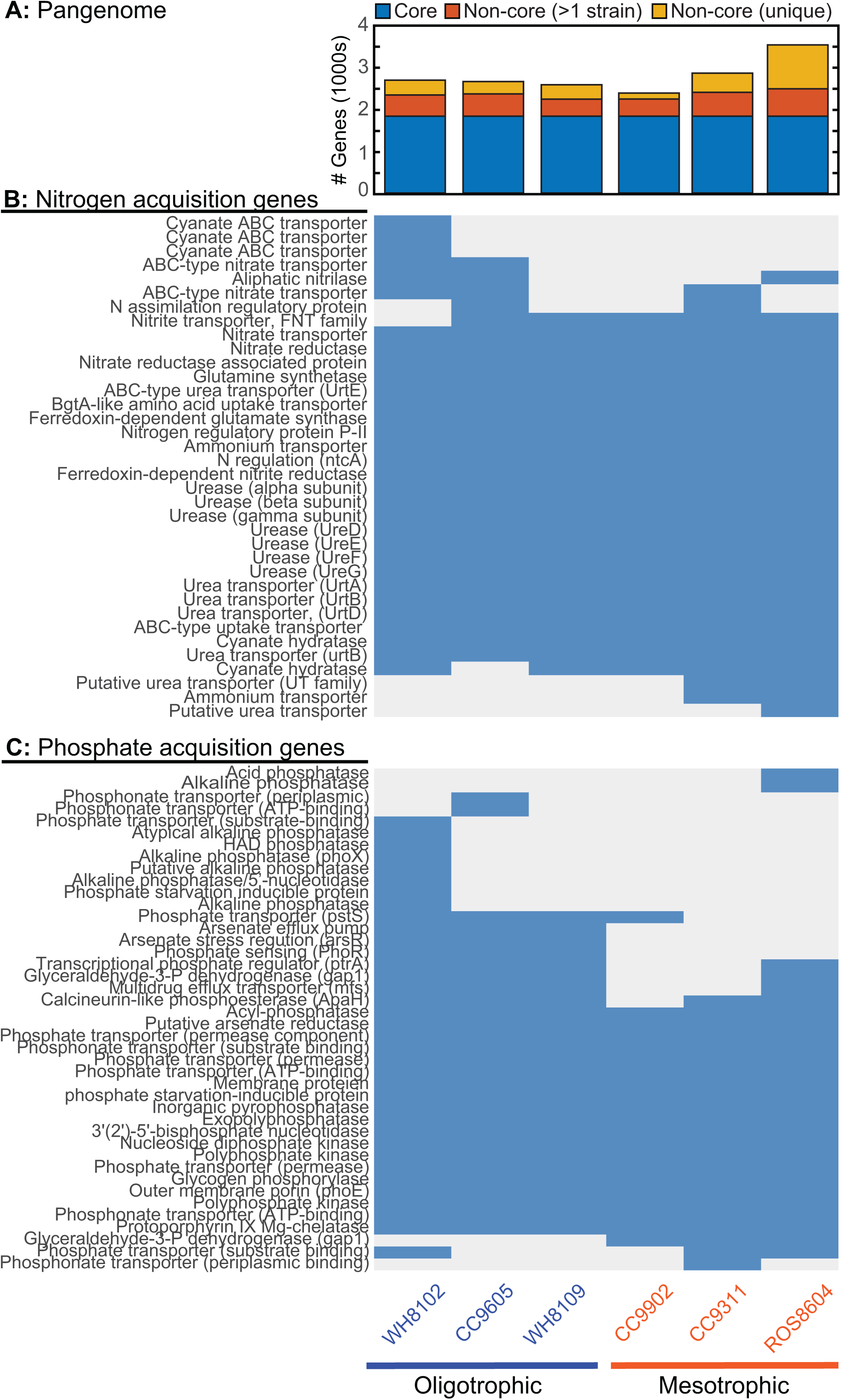
*Synechococcus* pan-genome. **A.** Genes classified into core (present in all six genomes) and non-core including both shared (>1 genome) and strain-specific (uniquely present in 1 strain). **B.** Genome presence of N acquisition genes. **C.** Genome presence of P acquisition genes.

**Table 1:**
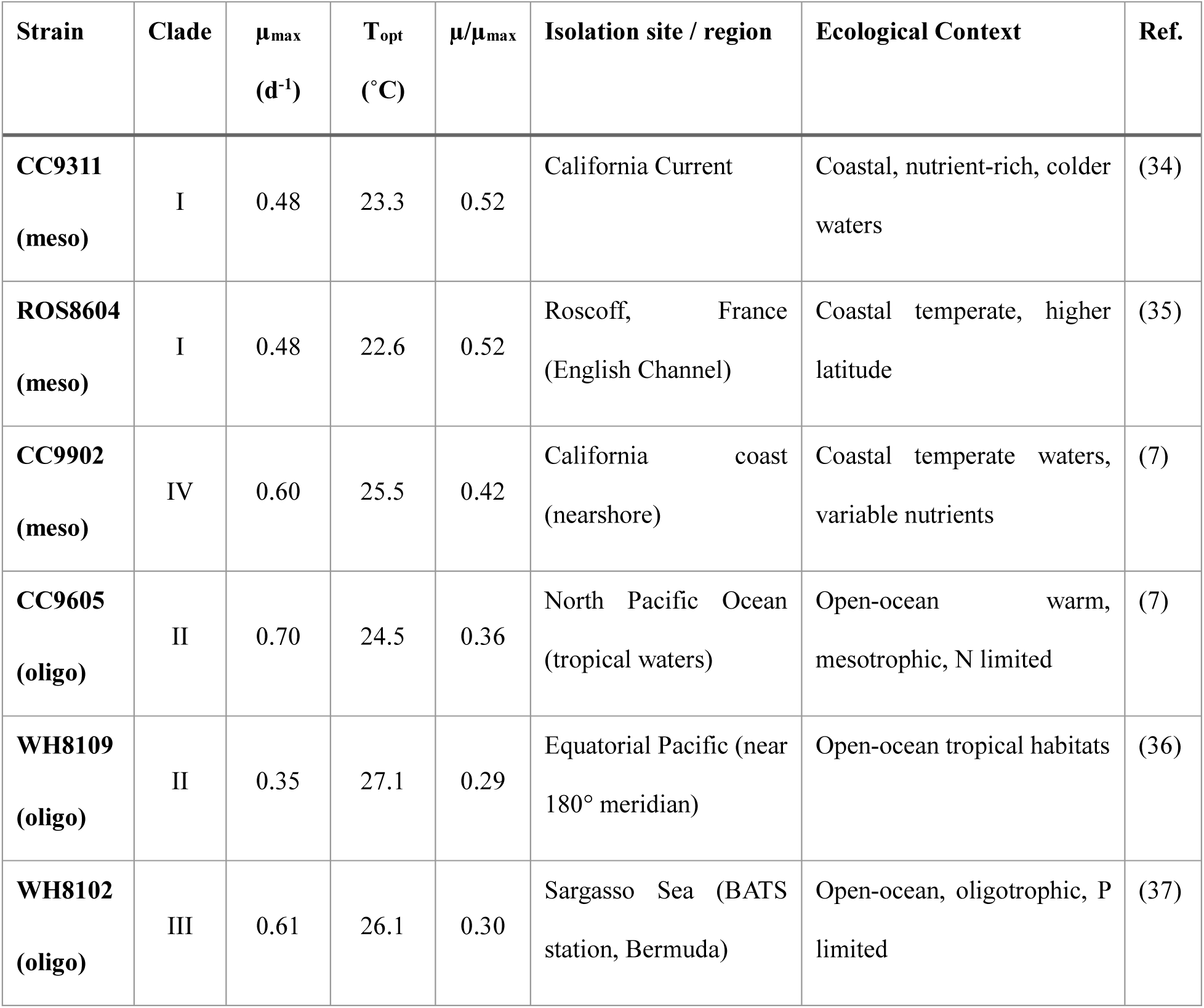
Summary of strains used in this study.

*Synechococcus* display significant cellular elemental plasticity to N vs. P stress. We grew the six strains under balanced growth in chemostats either N (*N:P_input_* = 1.7) or P stressed (*N:P_input_* = 80). Through monitoring cell counts and nutrient measurements, we observe that it took approximately thirty days for the chemostat cell cultures to reach steady state (Fig. S2). The limiting nutrient is consumed below detection (Fig. S3). Each lineage has a distinct cell size with ROS8604_meso_ being largest (diameter = 1.2 µm) and WH8109_oligo_ and CC9902 _meso_ being smallest (diameter = 0.7 µm) (Fig. 2A). Generally, cell size and the cell quotas positively correlate closely (Fig. 2H and Fig. S4) (20, 23). Cells are significantly larger under P compared to N stress, leading to a larger carbon quota by an average of 23 fg C under P stress (Fig. 2B). This cell enlargement also correlates to significantly higher N quotas under P stress except for WH8109_oligo_ that has lower N quotas under P stress (Fig. 2C). The P quota shows the opposite trend. Despite cells being larger under P stress, the P quota is significantly lower (Fig. 2D). There is a parallel response of the cellular elemental stoichiometry to N and P stress. The change in the cellular P quota strongly influences C:P and N:P, with both ratios significantly higher under P stress (Fig. 2 F,G). C:N response is more variable, including high variance within each strain and treatment and thus some but not all strains have higher C:N under N stress (Fig. 2E). We also examined if variation C:N:P was linked to how close a strain was growing to its optimal conditions (i.e., µ/µ_max_) (24) but observed no significant relationship. As seen in other phytoplankton studies (15, 19), most *Synechococcus* lineages respond by reducing demand and associated cell quotas of the limiting element, leading to elevated carbon:nutrient ratios when stressed.

**Figure 2.**
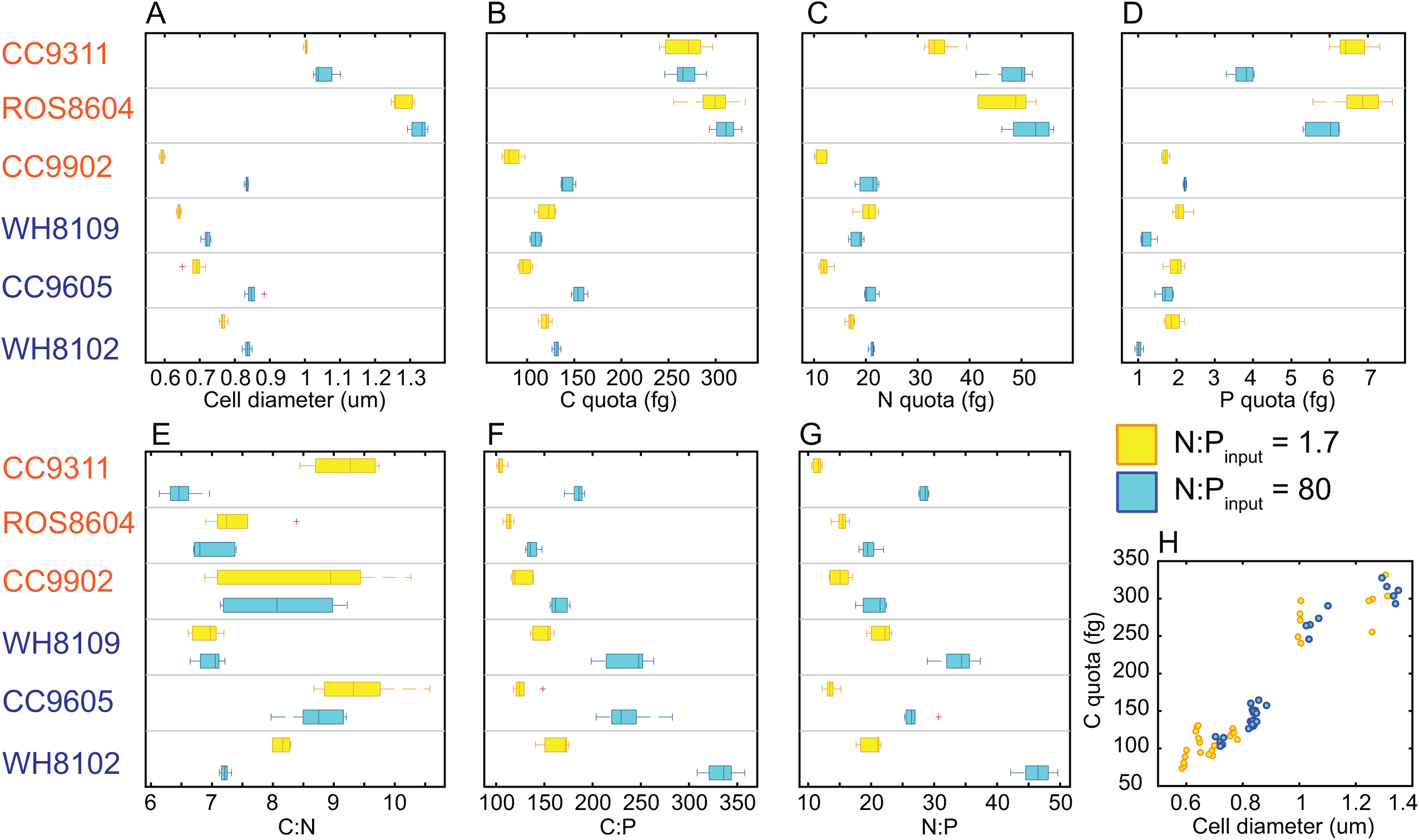
Strain-specific elemental response to nutrient stress. **A.** cell diameter based on an empirically derived conversion from FSCH and cell diameter (Fig. S1), **B.** cell carbon quota, **C.** cell nitrogen quota, **D.** cell phosphorus quota, **E.** carbon-to-nitrogen ratio (mol:mol), **F.** carbon-to-phosphorus ratio, **G,** nitrogen-to-phosphorus ratio. Boxes represent the 25-75 percentile, whiskers represent 99.3 percent coverage (standard Matlab settings), and n = 5-6. See Table S5 for full statistics. **H.** Significant relationship between cell diameter and biomass (*R_Pearson_*= 0.95, *p* < 1E-30). See Figure S3 for relationship between cell size and all cell quotas. Strains are grouped by environment, where strains in orange are isolated from mesotrophic and blue are from oligotrophic environments. There are six strains, two treatments, and 5-6 technical replicates and thus n_samples_ = 62.

In addition to the expected general effect of nutrient stress, the comparison also reveals lineage-specific resource demands. First, the mean elemental cell quotas and stoichiometry vary significantly across strains (Table S5, Fig. 2). As expected, larger cells (e.g., ROS8604_meso_) generally also have larger cell quotas (Fig. 2H). However, the mean cellular elemental stoichiometries vary without a clear size dependence (Figure 2). While the average C:N ratio is highest in the oligotrophic ocean strain CC9605_oligo_, some of the mesotrophic strains (e.g., CC9902_meso_) also have high C:N. The mean C:P and N:P correspond to the environmental origin, whereby open ocean, low nutrient strains (i.e., WH8102_oligo_, WH8109_oligo_, and CC9605_oligo_) on average have higher C:P and N:P than the mesotrophic strains. In addition to the distinct strain variation in mean elemental quotas and ratios, there are also clear differences in the elemental plasticity in response to a changing nutrient supply. C:N changes strongly in CC9311_meso_ and WH8102_oligo_, whereas the responses are muted in the remaining strains. A strain’s N:P and C:P plasticity generally aligns with habitat – oligo- vs. mesotrophic environments. The plasticity is highest in WH8102_oligo_, where the C:P ratio shifts by 175 (mol/mol) and N:P by 25 (mol/mol) from N to P stress (Figure 2F, G). In contrast, several mesotrophic strains display muted responses. The only exception is CC9311_meso_, which in addition to C:N, also shows strong changes in C:P and N:P. In sum, both the mean level and plastic responses of C:P and N:P are generally higher in oligo- vs. mesotrophic strains, whereas C:N varies in a less predictable way.

We observe a strong *N:P_input_* effect on the proteome within each strain (Fig. S5). Between 57 and 239 proteins are significantly differentially regulated within each strain (Fig. S5). Under N stress, many known nitrogen assimilation proteins are upregulated. These include proteins responsible for uptake and processing of ammonium, urea, nitrite, nitrate and cyanate. Many of these proteins display high peak area intensities (PAI) and thus make big contributions to the total protein biomass. Regulatory proteins including PII and NtcA are also differentially regulated in many strains. Under P stress, many known phosphorus assimilation proteins are induced. Proteins involved in the uptake of inorganic phosphate are highly expressed and constitute a large proportion of the total PAI. Differentially expressed proteins include an outer membrane porin (PhoE) and the ABC transporter system (PstABCS). Proteins annotated as phosphonate transporters are also differentially expressed. Hence, the proteomes in all strains are strongly impacted by shifts between N and P stress.

We next analyze the *Synechococcus* pan-proteome response and find that strain identity dominates proteome differences. First, we observe a significant strain effect on overall protein expression (Fig. 3A,C, Table S6). Despite the controlled culture conditions and strong physiological impact of N vs. P stress, the pan-proteome cluster distinctly by strain and much less by *N:P_input_* treatment (Fig. 3A). Strain identity explains 64 to 97% of total pan-proteome variance depending on normalization technique, whereas treatment (*N:P_input_*) contributes less than 6% (Fig. 3C, ‘All’). We expect this pan-proteome separation to be attributed to the many non-core genes. When we did a proteome ordination of only core genes, we still observe a similar strong separation in expression among strains (Fig. 3C, ‘Core’). Hence, the genome variance strongly affects the stress response of both core and non-core genes.

**Figure 3.**
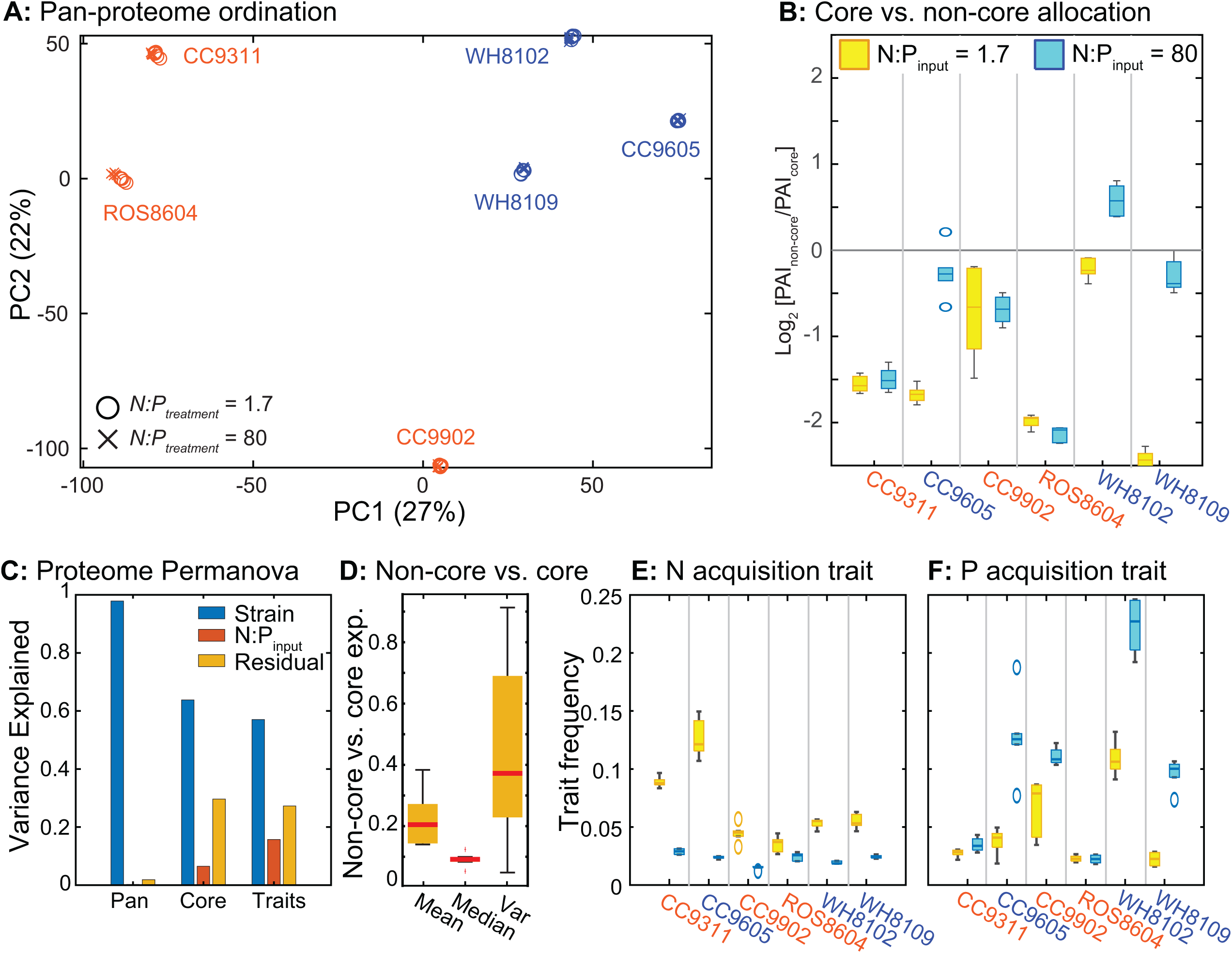
Pan-proteome response to nutrient stress. **A.** Strong strain effect on the pan-proteome response to N vs. P stress using principal component analysis (see Figure S5 for ordination of core protein expression profiles). **B.** Summed expression from core vs. non-core proteins showing that under most conditions core proteins dominate. However, non-core proteins are generally more expressed under P stress. **C.** Expression variance by strain identity vs. treatment using Permanova of pan-proteome (All), core proteins (Core), or 17 traits (Traits), see Table S1 for specific genes. **D.** Properties of Non-core vs core genome expression (log10 transformed) including significant differences (t-test, p < 0.05) in mean level, median level, and variance. **E, F.** Trait frequency by strain and treatment. Frequency is estimated as summed PAI for proteins within a specific trait divided by total PAI for that sample. Strain names are color-coded according to isolation environment (blue = oligotrophic, orange = mesotrophic).

Next, we detect a clear link between the pan-proteome response and controls on cellular C:N:P (Fig. 4A). First, we filtered out the strain-specific effect on protein expression (see Methods) so that subsequent analyses focused on proteomic responses shared across strains. We then asked whether coordinated changes in the proteome could explain the observed variation in cellular C:N:P. To address this question, we used partial least-squares (PLS) regression with the strain-normalized pan-proteome as the predictor matrix and cellular C:N:P as the response variables. PLS identifies combinations of proteins whose coordinated expression best explains variation in cellular stoichiometry, rather than testing proteins individually, thereby linking the global proteomic response to cellular elemental composition. For predicting cellular N:P, the best ‘variable-importance-in-projection’ (VIP) scores (Fig. 4C) belong to known nutrient stress response genes (Fig. 4D) with high induction levels (Fig. 4B). High cellular N:P is aligned with upregulation of some core proteins encoding for phosphate and phosphonate transport. Other core proteins with high VIP scores are involved in the oxidative pentose phosphate pathway, ATP production, peptidoglycan, and glycogen metabolism. Several stress response proteins are also associated with high N:P including GroEL and HtpG. Hence, a portion of the P stress response seen previously in strain WH8102_oligo_ is shared across this wider set of strains and encoded in core genome (15). Low cellular N:P is also aligned with upregulation of known core proteins involved in N stress regulation (e.g., NtcA and GlnB) as well as ammonium, urea, nitrite, nitrate, and cyanate assimilation. As seen previously in individual strains, N stress induces the production of the N-free compatible solute glycosyl-glycerate (GGA) and a down-regulation of peptidoglycan across *Synechococcus* (15). Finally, elevated PAI for ribosomal proteins align with low cellular N:P (Fig. 4). Again, these responses have been detected previously, but we now see they are shared across diverse strains. The pan-proteome expression also correlates with shifts in C:N or C:P. For example, high C:N is associated with many N stress proteins and high C:P is similarly linked to many P stress proteins. Hence, variations in C:N:P are aligned with core-protein shifts encoding nutrient stress as well as other core metabolic reactions.

**Figure 4.**
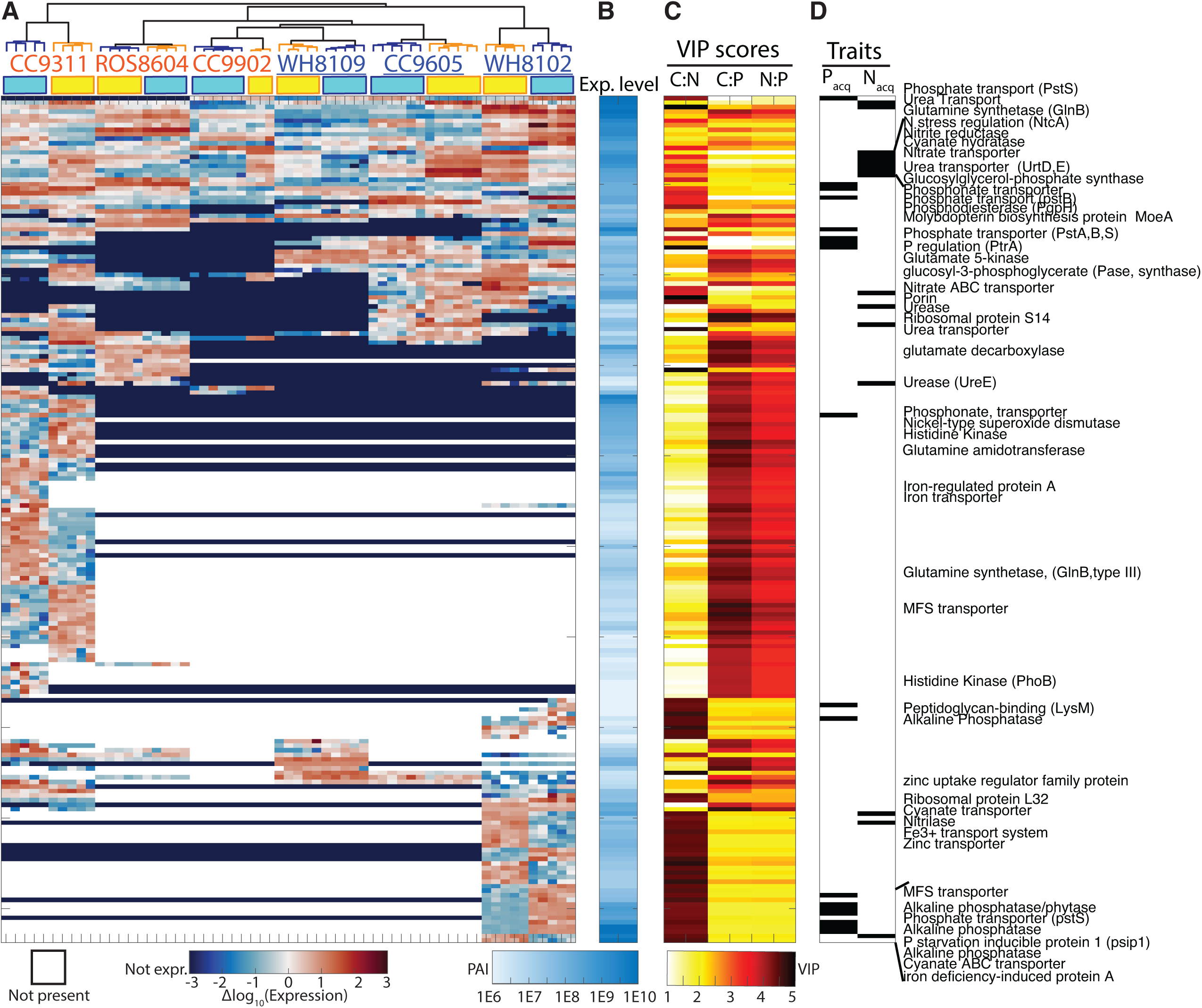
Linking pan-proteomic response and C:N:P regulation. **A:** Clustering of normalized expression of 187 proteins with VIP scores above 3.5. The plot is organized using 2-way hierarchical clustering with average linkage. White color represents an absent gene in that strain (i.e., non-core) and dark blue represents proteins with absent or very low expression. **B:** Mean expression level (PAI) of each selected protein across all treatments. **C:** VIP scores for predicting C:N, C:P, and N:P for each protein (see methods for how VIP scores are calculated). **D:** Nutrient acquisition trait membership of each protein (Table S1). VIP scores are identified using partial least-squares regression on the strain-normalized pan-proteome. Absence of genes in a strain (i.e., non-core genes) are represented with white color in panel 4A. Strain names are color-coded according to isolation environment (blue = oligotrophic, orange = mesotrophic).

The strongest response to *N:P_input_* is detected in non-core proteins. Compared to core proteins, expression of many non-core proteins with their variable presence among the strains, are significantly more sensitive to shift in *N:P_input_* (Fig. 3D). Indeed, both the absolute difference and variance in expression are elevated among non-core genes. Thus, non-core genes vary both among the strains and in expression level and thus central to the nutrient stress response. Furthermore, the expression of many non-core proteins is strongly aligned with variation in cell biomass composition (Fig. 4). High cellular N:P is associated with a strong induction of known P stress proteins including diverse alkaline phosphatases, diphosphatases, and P starvation regulators (e.g, PtrA). These non-core genes have high expression (PAI) and thus make important quantitative contributions to biomass allocation (Fig. 4B). Several P stress proteins previously identified in *Prochlorococcus* (11) are also tied to high cellular N:P including a multidrug efflux transporter (Mfs) and glycerol-3-phosphate dehydrogenase (Gap1). Non-core functions with high VIP score and aligned with low cellular N:P are linked to nitrile (i.e., triple-bonded N) utilization as well as an outer membrane porin. As far as we know, there is no porin in *Synechococcus* dedicated to N-compounds, so the role of this porin found in half the strains is unknown but likely important for N assimilation. We also observe multiple genes responsible for metal uptake. This expression profile indicates a link between N and metal nutrient stress, possible to use as co-factors in key proteins. Hence, non-core proteins involved in P and N stress responses are significant mechanisms for cellular variation in C:N:P.

The pan-proteome shifts translate into distinct trait expressions. Trait expression was calculated as the summed expression (as reflected by PAI) of all genes associated with a specific function. The largest trait investment is producing phycobilisomes (18% of total PAI, Fig. S7). Combined with photosynthetic electron transport, photosynthesis, and carbon fixation traits, investments in a phototrophic lifestyle constitute on average 32% of total PAI. Other abundant traits include solute production, peptidoglycan biosynthesis, stress response, and nutrient uptake, and collectively these few traits explain the majority of total protein expression (Fig. S7). Similar to the pan-proteome, strain identity is also the dominant contributor to trait expression, whereas treatment explains less than 3% of the variance (Fig. 3C). Thus, the shift from N to P limitation results in relatively modest changes in overall trait expression. However, nutrient acquisition representing the primary exception (Fig. 3D–F). Expression of the N acquisition trait triples from 2.4% of total PAI under P limitation to 7.2% under N limitation (Fig. 3E). Likewise, P acquisition increases from 2.3% to 8.7% of total PAI when cells shift from N to P limitation.

Expression changes in nutrient stress traits significantly parallel shifts in cellular elemental stoichiometry (Fig. 5). The magnitude of these responses differs markedly among strains and broadly follows their ecological origins (Table 1). WH8102_oligo_, an oligotrophic strain isolated from the strongly P-limited Sargasso Sea (25), exhibits the largest increase in P acquisition, reaching more than 20% of total PAI under P limitation, together with the strongest increases in C:P and N:P (Fig. 5A, B). The closely related Sargasso Sea strain WH8109_oligo_ shows a similar but more moderate response. In contrast, the oligotrophic North Pacific strain CC9605_oligo_ displays a comparatively stronger response to N limitation than to P limitation, consistent with its origin from an N-limited region (10, 26). Among the mesotrophic strains, CC9311_meso_ - also from the N limited eastern Pacific but affiliated with a mesotrophic ecotype - exhibits substantial plasticity in N acquisition and C:N (Fig. 5C). In contrast, another mesotrophic strain with a larger cell size, ROS8604_meso_, only shows only limited changes in both nutrient-acquisition traits and cellular C:N. Together, these observations demonstrate that variation in nutrient-acquisition trait expression is closely associated with differences in cellular stoichiometric plasticity across diverse *Synechococcus* strains.

**Fig. 5.**
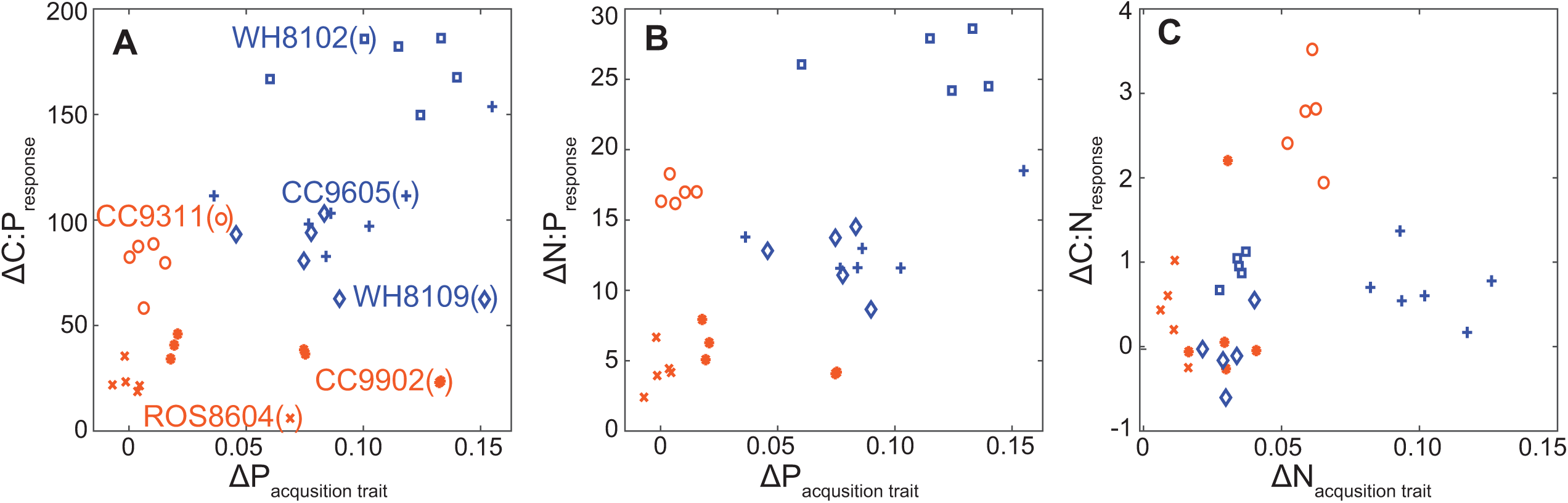
Relationship between treatment induction of C:N:P and proteome trait expression. **A:** Treatment change in C:P (ΔC:P) vs. treatment change in proteome expression of P traits (ΔP_acquisition_) (*R_Pearson_* = 0.75, *p* < 1E-6). **B:** Treatment change in N:P (ΔN:P) vs. treatment change in proteome expression of P traits (ΔP_acquisition_) (*R_Pearson_* = 0.54, *p* < 5E-3). Treatment change in C:N (C:N) vs. treatment change in proteome expression of N traits (ΔN_acquisition_) (*R_Pearson_* = 0.25, *p* > 5E-2). Strain names are color-coded according to isolation environment (blue = oligotrophic, orange = mesotrophic).

## Discussion

Our results demonstrate that local adaptation and associated strain diversity translates into predictable variation in physiological trait expression, which in turn connects to cellular C:N:P plasticity in a key marine phytoplankton lineage *Synechococcus*. Rather than responding uniformly to nutrient limitation, strains differed markedly in their ability to deploy nutrient-acquisition traits, and these differences closely paralleled shifts in cellular elemental composition. Oligotrophic lineages generally exhibited the greatest physiological flexibility, whereas mesotrophic strains showed more constrained responses, suggesting that adaptation to contrasting nutrient regimes has shaped the capacity for stoichiometric plasticity. At the same time, differences among strains occupying broadly similar environments indicate that clade affiliation further modifies these responses. Together, our findings support a mechanistic model in which ecological adaptation defines the repertoire of nutrient-response traits which in turn controls proteomic plasticity and ultimately align genome diversity to elemental stoichiometry and hence an important coupling between the marine nutrient and carbon cycles.

Across studies, assimilation pathways emerge as a key component of the molecular nutrient stress response. The strong induction of nitrogen transporters (nitrate, urea, ammonium, cyanate) and regulators such as NtcA and GlnB under N limitation, and phosphate plus phosphonate transporters under P limitation, is consistent with earlier work in both *Synechococcus* and *Prochlorococcus* (27). Many nutrient stress functions including a diverse set of alkaline phosphatases are encoded in the non-core genome and are known to vary among strains (28), yet when present they show highly predictable regulation. This underscores how the pan-proteome both constrains and enables a shared response within each nutrient acquisition trait. While a pan-genome reflects a species’ collective genetic potential, the pan-proteome reflects the subset of those genes that are functionally deployed under the experimental and environmental conditions studied. Other metabolic adjustments show broad parallels with prior studies. Here, we see an upregulation of several putative N-saving strategies including N-free compatible solutes and reduced peptidoglycan synthesis (Fig. 4). Similarly, P limitation induces proteins in the oxidative pentose phosphate pathway and glycogen metabolism, consistent with redirection of carbon fluxes under low P (Fig. 4). Metal uptake genes, variably present across strains, are also linked to N stress responses, suggesting higher cofactor demand for nitrogen assimilation, for example Fe in nitrate reductase, Ni in urease and Fe or Co in nitrile hydratase enzymes. In sum, the conserved induction of nutrient assimilation traits reflects a shared molecular core, whereas variation in non-core metabolic adjustments highlights how pan-proteome diversity modulates the breadth of physiological plasticity under different nutrient stress types.

The exact molecular mechanisms controlling cellar elemental composition are less clear. Ribosome concentration is often invoked as the primary driver of C:N:P both marine and non-marine ecosystems (1, 29). In support, induction of ribosomal proteins is aligned with low N:P. This was unexpected due to the controlled growth rate in our chemostat but implies an important role of ribosomes in regulating the observed C:P and N:P changes. We also observe strong induction of many uptake proteins under especially P stress. Concurrently, cells are larger. Hence, part of the higher C:P and N:P under P stress can be due to this strong protein induction, but we lack cellular protein concentration measurements to fully evaluate this hypothesis. The P quota is also lower which past measurements have been shown to in part due to lower phospholipids (30). However, we suspect that other P- or N-containing molecules not measured here are responding too. It is currently challenging to accurately measure the concentration of diverse cellular macromolecules and metabolites simultaneously. Hence, directly linking the proteome changes into a detailed chemical remodeling of the cell is difficult.

Together, our findings reveal that the genomic diversity and pan-proteome regulation of *Synechococcus* define the mechanistic basis of phytoplankton buffering capacity to future stratification and nutrient stress. Just as biodiversity broadens the thermal performance range of marine primary producers (31, 32), genomic and proteomic variation among lineages expand the stoichiometric response space of phytoplankton to nutrient stress. The capacity of populations to maintain balanced elemental composition under shifting nutrient regimes emerges not from a single genotype’s plasticity, but from the collective flexibility encoded within the pan-genome. This alignment between evolutionary adaptation and physiological acclimation enables communities to modulate their cellular stoichiometry beyond the capability of single lineage. As a result, large-scale patterns of oceanic C:N:P can remain predictable from ambient nutrient conditions (33), not because stoichiometry is solely driven by biodiversity or physiology, but because genomic potential and resultant proteome composition are selected from a large pool of global biodiversity to act in concert to buffer ecosystem responses. Such a genome-by-environment framework thus provides a simple model for linking complex biological processes to the large-scale regulation of nutrient stress, ecosystem stoichiometry, and ocean biogeochemical cycles.

## Materials and Methods

### Experimental design and culturing conditions

This study builds directly on a methodological framework previously published (15, 20), but with some modifications to allow for robust growth across all isolates. Specifically, in this study we (i) examined five additional *Synechococcus* strains plus the original analysis of strain WH8102_oligo_, and (ii) supplied nitrogen as an even mixture of nitrate (NO₃⁻) and urea as some strains struggled using solely nitrate.

As done in the past, we grew *Synechococcus* strains CC9311_meso_, CC9605_oligo_, CC9902_meso_, WH8109_oligo_ (RCC2033), and ROS8604_meso_ in continuous chemostat cultures (Table 1). Each strain represents a lineage adapted to distinct nutrient regimes in the global ocean. Cultures were maintained in 2 L polycarbonate vessels at 22 °C with gentle stirring and aeration using 0.2 μm-filtered air. Illumination was provided by cool white fluorescent lamps between 75 and 125 μmol photons m⁻² s⁻¹ under a 12 h light : 12 h dark cycle.

Artificial seawater medium was prepared following the formulation in Garcia et al. (2024) with phosphorus was supplied as phosphate (PO₄³⁻) but nitrogen was supplied as a 1:1 molar mixture of nitrate and urea (as some strains struggled with growing on nitrate as sole N source). To impose nutrient limitation, input N:P ratios (*N:P_input_*) were 1.7 (N-limited) or 80 (P-limited). Chemostats were run at a dilution rate of 0.25 d⁻¹ to maintain balanced exponential growth. The only exception was WH8109_oligo_ which was run at 0.1 d^-1^ to account for this overall slower growing isolate. We also included previously published observations for WH8102_oligo_ for comparison, where the temperature was 2°C higher at 24°C, nitrate was the sole N source, and a dilution rate of 0.18 d⁻¹. The strains also vary between 0.35 and 0.70 in the maximum growth rate. There was also a range in the temperature optima with some growing close to the opima whereas others grew a few degrees below the optima (Table 1). Hence, the experienced growth conditions are very similar but not completely identical due to individual biological behavior among the strains.

### Monitoring and sample collection

Culture dynamics were monitored by flow cytometry (Novocyte 1000, Acea Biosciences) to measure cell density and forward scatter (FSCH, proxy for cell size). Here, we used a 488 nm laser for excitation and monitored forward scatter as well as orange (575 nm) and red fluorescence (640 nm). Steady-state was verified by stable cell densities, nutrient concentrations below detection, and reproducible cellular quotas across replicate samples. For elemental analyses, replicate culture samples were collected during the mid-light period after at least five residence times to ensure acclimation. Particulate organic carbon (POC) and nitrogen (PON) were collected on pre-combusted GF/F filters (Whatman), dried, and analyzed with a Flash EA1112 elemental analyzer. Particulate organic phosphorus (POP) was collected on separate pre-combusted GF/F filters, dried with magnesium sulfate, digested at 450 °C for 2 h, hydrolyzed with HCl, incubated 2:5:1:2 parts ammonium molybdate tetrahydrate, 5 N sulfuric acid, potassium antimonyl tartrate, and ascorbic acid for 30 min, and measured colorimetrically at 885 nm. Nutrient concentrations were measured at the Marine Science Institute, University of California, Santa Barbara. We monitored the culture purity by staining cells with SYBRGreen and counting them with the flow cytometry (excitation 488 nm/emission 525/45 nm). The median number of stained particles without pigment fluorescence represented less than 5% of all cells including achloretic *Synechococcus* lacking pigment fluorescence.

### Pan-genome analysis and trait aggregation

We calculated full-genome amino-acid identities (AAI) using EzAAI (38). We compared AAI using hierarchical clustering with average linkage. Genes and associated proteins were assigned to orthologous groups based on CyanoRak v2.1 annotations (39). Core proteins were defined as present in all six strains whereas non-core includes shared (present in more than one but not all) and strain-specific being unique to one strain. The pan-proteome spans both core and non-core proteins. The genome content of each strain was compared using the Jaccard index and summarized using hierarchical clustering with average linkage (Matlab ‘clustergram’). Proteins were aggregated into 17 functional trait categories (Table S1) based on CyanoRak annotations and prior trait groupings (15). The trait frequency was calculated as the summed sample PAI by trait normalized to total PAI.

### Protein extraction and proteomic workflow

Cells were collected on 0.2 μm polycarbonate filters (47 mm), pelleted by centrifugation at 21,000 g for 3 min, flash frozen in liquid nitrogen, and stored at −80 °C until extraction. Proteins were extracted in 50 mM HEPES (pH 8.5) with 1% SDS by heating at 95 °C for 10 min followed by 30 min of shaking at room temperature. Lysates were treated with Benzonase nuclease (Novagen) to degrade nucleic acids, reduced with 200 mM dithiothreitol (DTT), and alkylated with 400 mM iodoacetamide. Protein extracts were purified using a modified SP3 (Single Pot 3) method (40) using SpeedBead magnetic particles (20 μg/μl) added to 400 μl of extracted protein sample, acidified with formic acid (pH of 2–3) and washed with ethanol and acetonitrile using a magnetic rack, quantified using a bicinchoninic acid (BCA) assay, and digested overnight with trypsin (Promega). Peptides were desalted, concentrated, and analyzed by data-independent acquisition mass spectrometry (DIA-MS) on a Q-Exactive mass spectrometer (Thermo Fisher) coupled to a Michrom Advance HPLC system. Peptides were separated on a C18 column (Reprosil-Gold, Dr. Maisch GmbH) using a 200 min acetonitrile gradient.

### Proteomic data processing

Raw DIA data were converted to mzML format and processed using Scaffold DIA (Proteome Software) and EncyclopeDIA. Searches were conducted against proteomes for each *Synechococcus* strain (Table 1) and peak area intensities (PAI) were reported. We subsequently normalized the proteome PAI data by strain by first logarithmically transforming followed by a quantile normalization (by strain, done in Scaffold DIA). We subsequently scaled all strains by their median intensity (Table S2). We also considered PAI observations without log-transformation (Table S3). We treated each proteome for missing observations. *First*, we evaluated if a protein was undetected in more than one sample for both treatments (i.e., *N:P_input_*) within a strain. If so, this protein was removed from this specific strain. *Second*, if a protein was completely absent in one treatment but present in nearly all samples from the other treatment (could be missing in one sample), then we replaced the absent treatment values with the minimum value PAI value for that strain in order to avoid the presence of zero values that disallow analyses involving log transformations. We interpret this profile as representing proteins absent in one treatment but upregulated in the other. Hence, our approach is designed to allow for this protein to be included in subsequent analyses. *Third*, if a protein is missing in one sample within a treatment, we replace the missing value with the median of the remaining observations within treatment and protein. We used a student’s t-test [Matlab: mattest(X_N:P=1.7,_, X_N:P=80,_, ‘Vartype’, ’equal’)] followed by a Benjamin-Hochberg procedure [Matlab: mafdr(ttest, ‘BHFDR’, true] to calculate the False Discovery Rate (FDR) and find significant treatment (*N:P_input_*) effects on protein expressions within each strain (Table S4). The proteome regulation in each strain was displayed using a volcano plot (colored dots in Fig. S5, FDR cutoff = 0.05 and a doubling in expression).

### Pan-proteome analysis

2-way differences (strain x treatment) in multivariate pan-proteome and trait expressions were assessed using principal component analysis (Matlab: PCA with Euclidian distance) and Permanova (R w. ‘vegan’ package: adonis2 w. Euclidian distance) (41). These analyses were done with (*norm*) or without (*obs*) logarithmic normalization for proteome expressions - although there was little difference in the outcome. Partial least-squares (PLS) regression was performed using the measured abundance of all proteins as predictor variables (X matrix) and the measured cellular elemental stoichiometry (C:N:P) of each biological replicate as the response variables (Y matrix) (42). Unlike separate regressions for individual proteins, PLS simultaneously models all proteins to identify latent variables that maximize the covariance between the proteome and elemental stoichiometry. Variable Importance in Projection (VIP) scores quantify the contribution of each protein to the predictive latent variables, with larger values indicating proteins that contribute more strongly to explaining variation in cellular stoichiometry. First, we removed the strain effect for each protein expression. To achieve this, we calculated the mean expression for the protein by strain and subtracted this mean strain-specific value from each expression value. Second, we took the zscore for each strain to achieve numerical stability. Third, we estimated the optimal number of PLS model components (N_comp_; where N_comp_ for C:N = 1, C:P = 4, and N:P = 4) by leaving-one-strain-out (LOSO) and minimize the root mean square error (RMSE). We then estimated the optimal PLS regression model [Matlab: plsregress(X,Y,N_comp_)] and calculated VIP (Variable Importance in Projection) scores:

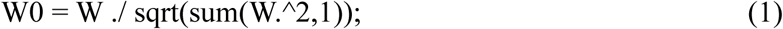

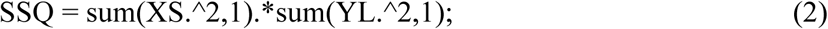

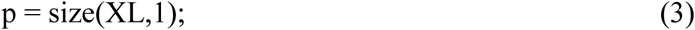

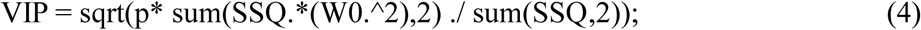

W is the weights, XS is the predictor scores, XL is the predictor loadings, and YL is the response loadings. These are all outputs from the PLS regression. Another output (beta) is the correlation between each protein expression and response variable. We combined VIP scores and beta to evaluate proteins across the pan-genome corresponding to high vs. low stoichiometry ratios.

## Acknowledgements

We thank Jennifer Martiny for many helpful suggestions for this study. We want to acknowledge the National Science Foundation for supporting this work (IOS-2137339, OCE-2135035, and OCE-2517928). The authors declare that they have no conflicts of interest regarding the publication of this paper.

## Data Availability

Biogeochemical chemostat data is present in Table S7. Proteome data can be found in the ‘PRIDE’ database (https://www.ebi.ac.uk/pride) under accession number PXD072227. The sample list with detail accession numbers is present in Table S8. All processed data and statistical analyses are present in Table S1 to S6.

## Supplementary Material

Table S1 – S8

Figure S1 – S7

## References

1. Moreno AR, Martiny AC. 2018. Ecological Stoichiometry of Ocean Plankton. Annu Rev Mar Sci 10:43–69.

2. Moore JK, Fu W, Primeau F, Britten GL, Lindsay K, Long M, Doney SC, Mahowald N, Hoffman F, Randerson JT. 2018. Sustained climate warming drives declining marine biological productivity. Science 359:1139–1143.

3. Geider RJ, La Roche J. 2002. Redfield revisited: variability of C : N : P in marine microalgae and its biochemical basis. Eur J Phycol 37:1–17.

4. Teng Y-C, Primeau FW, Moore JK, Lomas MW, Martiny AC. 2014. Global-scale variations of the ratios of carbon to phosphorus in exported marine organic matter. Nat Geosci 7:895–898.

5. Kwon EY, Sreeush MG, Timmermann A, Karl DM, Church MJ, Lee S-S, Yamaguchi R. 2022. Nutrient uptake plasticity in phytoplankton sustains future ocean net primary production. Sci Adv 8:eadd2475.

6. Kettler GC, Martiny AC, Huang K, Zucker J, Coleman ML, Rodrigue S, Chen F, Lapidus A, Ferriera S, Johnson J, Steglich C, Church GM, Richardson P, Chisholm SW. 2007. Patterns and Implications of Gene Gain and Loss in the Evolution of *Prochlorococcus*. PLoS Genet 3:e231.

7. Dufresne A, Ostrowski M, Scanlan DJ, Garczarek L, Mazard S, Palenik BP, Paulsen IT, de Marsac NT, Wincker P, Dossat C, Ferriera S, Johnson J, Post AF, Hess WR, Partensky F. 2008. Unraveling the genomic mosaic of a ubiquitous genus of marine cyanobacteria. Genome Biol 9:R90.

8. Berube PM, Rasmussen A, Braakman R, Stepanauskas R, Chisholm SW. 2019. Emergence of trait variability through the lens of nitrogen assimilation in Prochlorococcus. eLife 8:e41043.

9. Mackey KRM, Post AF, McIlvin MR, Cutter G a, John SG, Saito M a. 2015. Divergent responses of Atlantic coastal and oceanic *Synechococcus* to iron limitation. Proc Natl Acad Sci 112:9944–9949.

10. Ustick LJ, Larkin AA, Garcia CA, Garcia NS, Brock ML, Lee JA, Wiseman NA, Moore JK, Martiny AC. 2021. Metagenomic analysis reveals global-scale patterns of ocean nutrient limitation. Science 372:287–291.

11. Martiny AC, Coleman ML, Chisholm SW. 2006. Phosphate acquisition genes in *Prochlorococcus* ecotypes: Evidence for genome-wide adaptation. Proc Natl Acad Sci U S A 103:12552–12557.

12. Tanioka T, Garcia CA, Larkin AA, Garcia NS, Fagan AJ, Martiny AC. 2022. Global patterns and predictors of C:N:P in marine ecosystems. Commun Earth Environ 3:1–9.

13. Garcia CA, Hagstrom GI, Larkin AA, Ustick LJ, Levin SA, Lomas MW, Martiny AC. 2020. Linking regional shifts in microbial genome adaptation with surface ocean biogeochemistry. Philos Trans R Soc B Biol Sci 375:20190254.

14. Hall EK, Bernhardt ES, Bier RL, Bradford MA, Boot CM, Cotner JB, del Giorgio PA, Evans SE, Graham EB, Jones SE, Lennon JT, Locey KJ, Nemergut D, Osborne BB, Rocca JD, Schimel JP, Waldrop MP, Wallenstein MD. 2018. Understanding how microbiomes influence the systems they inhabit. Nat Microbiol 3:977–982.

15. Garcia NS, Du M, Guindani M, McIlvin MR, Moran DM, Saito MA, Martiny AC. 2024. Proteome trait regulation of marine *Synechococcus* elemental stoichiometry under global change. ISME J 18:wrae046.

16. Tetu SG, Brahamsha B, Johnson DA, Tai V, Phillippy K, Palenik B, Paulsen IT. 2009. Microarray analysis of phosphate regulation in the marine cyanobacterium *Synechococcus* sp WH8102. ISME J 3:835–849.

17. Litchman E, Klausmeier CA. 2008. Trait-Based Community Ecology of Phytoplankton. Annu Rev Ecol Evol Syst 39:615–639.

18. Broadbent JA, Broszczak DA, Tennakoon IUK, Huygens F. 2016. Pan-proteomics, a concept for unifying quantitative proteome measurements when comparing closely-related bacterial strains. Expert Rev Proteomics 13:355–365.

19. Rhee GY. 1978. Effects of N-P atomic ratios and nitrate limitation on algal growth, cell composition, and nitrate uptake. Limnol Oceanogr 23:10–25.

20. Garcia NS, Bonachela JA, Martiny AC. 2016. Interactions between growth-dependent changes in cell size, nutrient supply and cellular elemental stoichiometry of marine *Synechococcus*. ISME J 10:2715–2724.

21. Zwirglmaier K, Jardillier L, Ostrowski M, Mazard S, Garczarek L, Vaulot D, Not F, Massana R, Ulloa O, Scanlan DJ. 2008. Global phylogeography of marine *Synechococcus* and *Prochlorococcus* reveals a distinct partitioning of lineages among oceanic biomes. Env Microbiol 10:147–161.

22. Doré H, Guyet U, Leconte J, Farrant GK, Alric B, Ratin M, Ostrowski M, Ferrieux M, Brillet-Guéguen L, Hoebeke M, Siltanen J, Le Corguillé G, Corre E, Wincker P, Scanlan DJ, Eveillard D, Partensky F, Garczarek L. 2023. Differential global distribution of marine picocyanobacteria gene clusters reveals distinct niche-related adaptive strategies. ISME J 17:720–732.

23. Harcourt R, Garcia NS, Martiny AC. 2024. Intraspecific trait variation modulates the temperature effect on elemental quotas and stoichiometry in marine *Synechococcus*. PLOS ONE 19:e0292337.

24. Godwin CM, Cotner JB. 2018. What intrinsic and extrinsic factors explain the stoichiometric diversity of aquatic heterotrophic bacteria? ISME J 12:598–609.

25. Martiny AC, Ustick L, A. Garcia C, Lomas MW. 2020. Genomic adaptation of marine phytoplankton populations regulates phosphate uptake. Limnol Oceanogr 65:S340–S350.

26. Gerace SD, Yu J, Moore JK, Martiny AC. 2025. Observed declines in upper ocean phosphate-to-nitrate availability. Proc Natl Acad Sci 122:e2411835122.

27. Scanlan DJ, Ostrowski M, Mazard S, Dufresne A, Garczarek L, Hess WR, Post AF, Hagemann M, Paulsen I, Partensky F. 2009. Ecological genomics of marine picocyanobacteria. Microbiol Mol Biol Rev 73:249–299.

28. Moore LR, Ostrowski M, Scanlan DJ, Feren K, Sweetsir T. 2005. Ecotypic variation in phosphorus acquisition mechanisms within marine picocyanobacteria. Aquat Microb Ecol 39:257–269.

29. Sterner RW, Elser JJ. 2002. Ecological stoichiometry: the biology of elements from molecules to the biosphere. Princeton University Press, Princeton, NJ.

30. Mouginot C, Zimmerman AE, Bonachela JA, Fredricks H, Allison SD, Van Mooy BAS, Martiny AC. 2015. Resource allocation by the marine cyanobacterium *Synechococcus* WH8102 in response to different nutrient supply ratios. Limnol Oceanogr 60:1634–1641.

31. Barton S, Jenkins J, Buckling A, Schaum CE, Smirnoff N, Raven JA, Yvon-Durocher G. 2020. Evolutionary temperature compensation of carbon fixation in marine phytoplankton. Ecol Lett 23:722–733 10.1111/ele.13469.

32. Martiny AC, Hagstrom GI, DeVries T, Letscher RT, Britten GL, Garcia CA, Galbraith E, Karl D, Levin SA, Lomas MW, Moreno AR, Talmy D, Wang W, Matsumoto K. 2022. Marine phytoplankton resilience may moderate oligotrophic ecosystem responses and biogeochemical feedbacks to climate change. Limnol Oceanogr 67:S378–S389.

33. Galbraith ED, Martiny AC. 2015. A simple nutrient-dependence mechanism for predicting the stoichiometry of marine ecosystems. Proc Natl Acad Sci U S A 112:8199–8204.

34. Palenik B, Ren QH, Dupont CL, Myers GS, Heidelberg JF, Badger JH, Madupu R, Nelson WC, Brinkac LM, Dodson RJ, Durkin AS, Daugherty SC, Sullivan SA, Khouri H, Mohamoud Y, Halpin R, Paulsen IT. 2006. Genome sequence of *Synechococcus* CC9311: Insights into adaptation to a coastal environment. Proc Natl Acad Sci U S A 103:13555– 13559.

35. Pittera J, Jouhet J, Breton S, Garczarek L, Partensky F, Maréchal É, Nguyen NA, Doré H, Ratin M, Pitt FD, Scanlan DJ, Six C. 2018. Thermoacclimation and genome adaptation of the membrane lipidome in marine *Synechococcus*. Environ Microbiol 20:612–631.

36. Waterbury JB, Valois FW. 1993. Resistance to co-occurring phages enables marine *Synechococcus* communities to to coexist with cyanophages abundant in seawater. Appl Environ Microbiol 59:3393–3399.

37. Palenik B, Brahamsha B, Larimer FW, Land M, Hauser L, Chain P, Lamerdin J, Regala W, Allen EE, McCarren J, Paulsen I, Dufresne A, Partensky F, Webb EA, Waterbury J. 2003. The genome of a motile marine *Synechococcus*. Nature 424:1037–1042.

38. Kim D, Park S, Chun J. 2021. Introducing EzAAI: a pipeline for high throughput calculations of prokaryotic average amino acid identity. J Microbiol 59:476–480.

39. Garczarek L, Guyet U, Doré H, Farrant GK, Hoebeke M, Brillet-Guéguen L, Bisch A, Ferrieux M, Siltanen J, Corre E, Le Corguillé G, Ratin M, Pitt FD, Ostrowski M, Conan M, Siegel A, Labadie K, Aury J-M, Wincker P, Scanlan DJ, Partensky F. 2021. Cyanorak v2.1: a scalable information system dedicated to the visualization and expert curation of marine and brackish picocyanobacteria genomes. Nucleic Acids Res 49:D667–D676.

40. Hughes CS, Moggridge S, Müller T, Sorensen PH, Morin GB, Krijgsveld J. 2019. Single-pot, solid-phase-enhanced sample preparation for proteomics experiments. Nat Protoc 14:68–85.

41. Oksanen J, Blanchet F, Kindt R, Legendre P, Minchin P, O’Hara R, Simpson G, Solymos P, Stevens M, Wagner H. 2013. vegan: Community Ecology Package. R package version 2.0–10.

42. Martens H, Næs T. 1992. Multivariate calibration. John Wiley & Sons.

